# Clear, extended field-of-view single-molecule imaging by highly inclined swept illumination

**DOI:** 10.1101/311753

**Authors:** Jialei Tang, Kyu Young Han

## Abstract

Highly inclined and laminated optical sheet (HILO) illumination facilitates high-contrast single-molecule imaging inside cells with a single objective lens. However, the beam thickness is strongly coupled to the illumination area, limiting its usage. Here, we present highly inclined swept tile illumination microscopy (HIST). By sweeping a thin HILO beam with confocal slit detection, HIST provides a 2-fold thinner illumination and >40-fold larger imaging area than conventional HILO microscopy, enabling 3D single-molecule imaging with high signal-to-background ratio. We demonstrate single-molecule mRNA imaging with a few probes or a single probe in cultured cells and mouse brain tissues.

## Main text

Single-molecule imaging is an indispensable tool in many biological studies, i.e., for revealing dynamics of biomolecules^1^, ultra-structures of sub-cellular components^2^, and spatial context of gene expression levels^3^. To visualize individual fluorescent molecules, it is critical to ensure a high signal to noise ratio. Although fluorescence signal can be amplified by multiple tagging systems^4^, their large size can potentially interfere with the activity of the target. Therefore, it is more desirable to decrease background level by minimizing unwanted out-of-focus fluorescent signal generated by the excitation beam in microscope systems. Total internal reflection fluorescence (TIRF) microscopy is widely used for single-molecule imaging^5^ due to the fact that it creates a shallow evanescent field, but TIRF or pseudo-TIRF^6^ is only capable of probing the bottommost ~1–2 μm of the cell. For higher imaging depth, highly inclined and laminated optical sheet (HILO) microscopy^7^ was suggested, where an incident beam refracts at glass/water interface with an angle slightly smaller than the critical angle, yielding a thin illumination with 3D sectioning capability. Unlike light-sheet based approaches^8–11^, HILO requires only a single high numerical aperture objective without an additional illuminator or reflector, and is compatible with typical sample chambers. These advantages have allowed HILO microscopy to be exploited for several research areas^12, 13^.

However, in HILO illumination, the beam thickness, *dz*, is closely related to the diameter of illumination beam, *R*, i.e. *dz* = *R*/tan(*θ*), where *θ* is the angle of transmitted beam (Fig. 1a)^7^. This means that a thin illumination unavoidably results in a small imaging area. For this reason, a previous study was only able to demonstrate <20 × 20 μm^2^ field-of-view (FOV) with approximately 6–7 μm beam thickness^7^, which makes it challenging to be used for imaging a large mammalian cell or multiple cells with high contrast. Here we overcome this limitation of HILO microscopy by sweeping a highly inclined beam elongated in one direction in conjunction with a confocal slit while maintaining advantages of HILO imaging.

**Figure 1.**
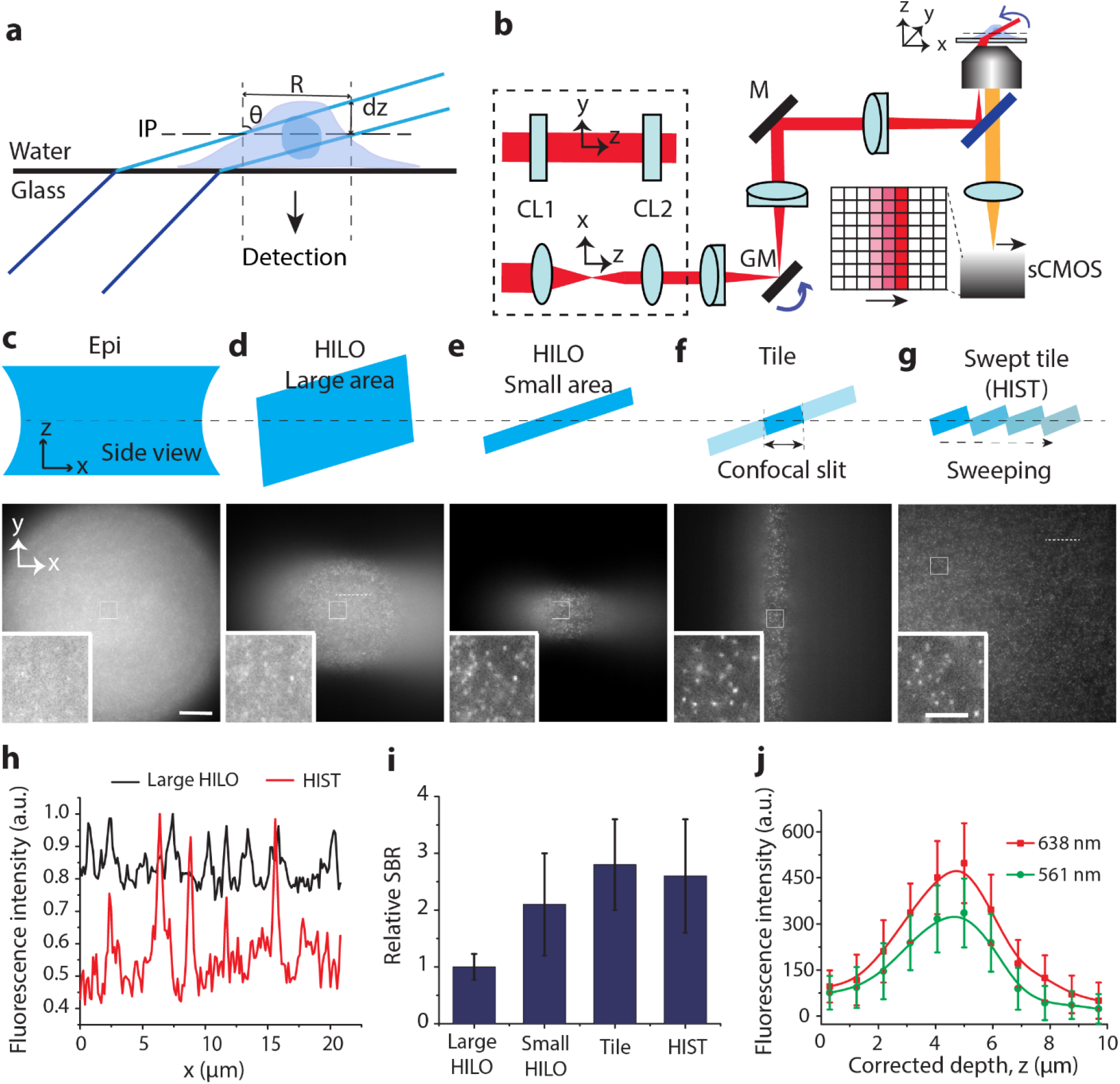
Highly inclined swept tiles (HIST) microscopy. (a) Highly inclined illumination at glass and water interface. *dz*, beam thickness; R, illumination width; IP, imaging plane. (b) Schematic of HIST microscopy. A tile beam is swept across the FOV by a galvo mirror and the emitted fluorescence is detected by an sCMOS camera with confocal slit detection. CL1–2, cylindrical lens; GM, galvo mirror; M, mirror. (c~g) Illumination schemes (top, *xz* cross-section) and single-molecule fluorescence images of Atto647N labeled DNA embedded in a hydrogel (bottom, *xy* cross-section) by epi (c), large area HILO (d), small area HILO (e), tile (f) and HIST illumination (g) at excitation wavelength of 638 nm. The images were taken 5 μm above the surface and the camera frame rate was 800 ms^−1^. The average illumination intensity was 30 W/cm^2^. Inset, zoom-in single-molecule images. All images are raw data. Scale bar, 20 μm and 1 μm (inset). (h) Line profiles of large area HILO and HIST illumination taken from (d) and (g) along white dashed lines. (i) Relative signal to background ratio (SBR) for each illumination method. More than 100 single-molecule spots were used for analysis. (j) Beam thickness of a highly inclined illumination beam (tile) with 638 nm and 561 nm excitation lights when 8x compression ratio is used. The beam thickness was 3.7 ± 0.3 μm and 3.6 ± 0.2 μm, respectively.

First, we created an elongated beam on the conjugated image plane using a pair of cylindrical lenses and sent it to our microscope with a high incidence angle like HILO illumination (Fig. 1b, Supplementary Fig. 1). For imaging, we prepared a 3D single-molecule hydrogel sample where each DNA probe was labeled with Atto647N and anchored to the hydrogel network via an acrydite moiety during gel polymerization (See Methods section). We imaged the sample at 5 μm above the surface (Fig. 1c-g). While in conventional HILO microscopy an iris controls the size of the illumination beam in both *x* and *y* directions^7^, our approach generates a tile-like beam elongated along the *y* axis on the sample plane, which is orthogonal to the direction of the illumination beam (Fig. 1f). This tile illumination increased the FOV from 20×20 μm^2^ to ~130×12 μm^2^. Importantly, it enabled the visualization of single-molecules with higher contrast than HILO illumination with *R* = 20 μm (Fig. 1e) because the elongation of the beam in the *y* axis does not affect the beam thickness. In contrast, single-molecule spots could hardly be detected with epi-illumination (Fig. 1c) or HILO illumination with a large beam size (Fig. 1d) due to the strong background, reconfirming that the image contrast of HILO illumination is highly dependent upon the beam size.

In order to extend the imaging area, we swept the tile along the *x*-axis by rotating a galvo mirror (Fig. 1b). However, the highly inclined beam generated out-of-focus background and blurred the image. To resolve this issue, we adopted confocal slit detection using a scientific complementary metal-oxide semiconductor (sCMOS) camera supporting a rolling shutter mode^14^. Synchronously sweeping the tile with the readout of the camera facilitated the rejection of background in real time without additional optical components^14^ (Supplementary Fig. 2). We have named our technique highly inclined swept tile (HIST) microscopy. Remarkably, we were able to clearly visualize single-molecules across ~130×130 μm^2^ FOV (Fig. 1g) which is more than 40 times larger than conventional HILO imaging. A line profile clearly showed the much improved signal-to-background ratio (SBR) of HIST microscopy and it was ~2.6-fold higher than HILO counterpart (Fig. 1h,i). The size of the total imaging area was decoupled from the beam thickness which solely depends on the width of tile, and thus it enabled a thinner illumination and larger FOV imaging.

We further characterized the performance of our method using 20-nm fluorescent beads embedded in 3D hydrogel with 638 nm excitation light. Firstly, we measured the effective illumination width and thickness of the tile at a compression ratio (*r*) of the beam that depends on the pair of cylindrical lenses. The measured width and thickness were ~10 μm and 3.7 ± 0.3 μm, respectively, at *r* = 8 when a ~80 μm long tile was used (Fig. 1j, Supplementary Fig. 3). If a lower compression ratio and/or a longer tile is used, the illumination beam becomes proportionally wider and thicker (Supplementary Fig. 3). For instance, SBR at *r* = 8 was 1.4-fold higher than SBR at *r* = 5 (Supplementary Fig. 4). The beam thickness with 561 nm light was 3.6 ± 0.2 μm (Fig. 1j) and a thin illumination was retained over a depth of >10 μm (Supplementary Fig. 5).

Since the tile is swept across a large FOV, the illumination angle (*θ*) was not constant along the *x* axis due to a slightly different refraction angle and aberration. It resulted in a rather elevated background level on the left side of HIST images (Fig. 1g). However, after background subtraction, a high contrast HIST image with a uniform background level was readily recovered (Supplementary Fig. 6, see Methods section). A fine adjustment by an additional galvo mirror instead of a manual mirror (M) in Fig. 1b will possibly help to keep the illumination angle constant during sweeping.

In addition, we measured the photobleaching kinetics by imaging *z*-stacks consisting of 40 images in order to check whether the 8 times higher instantaneous illumination intensity of HIST imaging may have adverse effects. However, the fluorescence intensity decay rate of HIST imaging was slower than that of epi-illumination at the same average excitation power, presumably due to less exposure to light (Supplementary Fig. 7).

To demonstrate potential applications of our method, we performed single-molecule RNA fluorescence *in situ* hybridization (smFISH) with a single probe^15, 16^ or a few probes^17^, which is critical for detecting single-nucleotide variants^15^ and rare transcriptional mutations. Figure 2a displays FISH images of *EEF2* (eukaryotic translation elongation factor 2) with four probes on A549 cells. The HIST image showed 1.5-fold higher SBR compared to its epi-illumination counterpart. The photobleaching step distribution revealed the actual number of probes (Supplementary Fig. 8). Further, we evaluated SBR of epi and HIST images with different numbers of FISH probes (*N* = 32, 24, 16, 12, 8, 4, 2, 1). When *N* ≤ 8, SBR of epi-illumination was not sufficient for robust detection and counting of molecules at our given experimental conditions, i.e. illumination intensity ~23 W/cm^2^ and exposure time 400 ms. In contrast, HIST microscopy showed high SBR (>2) even with a single probe (Fig. 2b, Supplementary Fig. 9).

**Figure 2.**
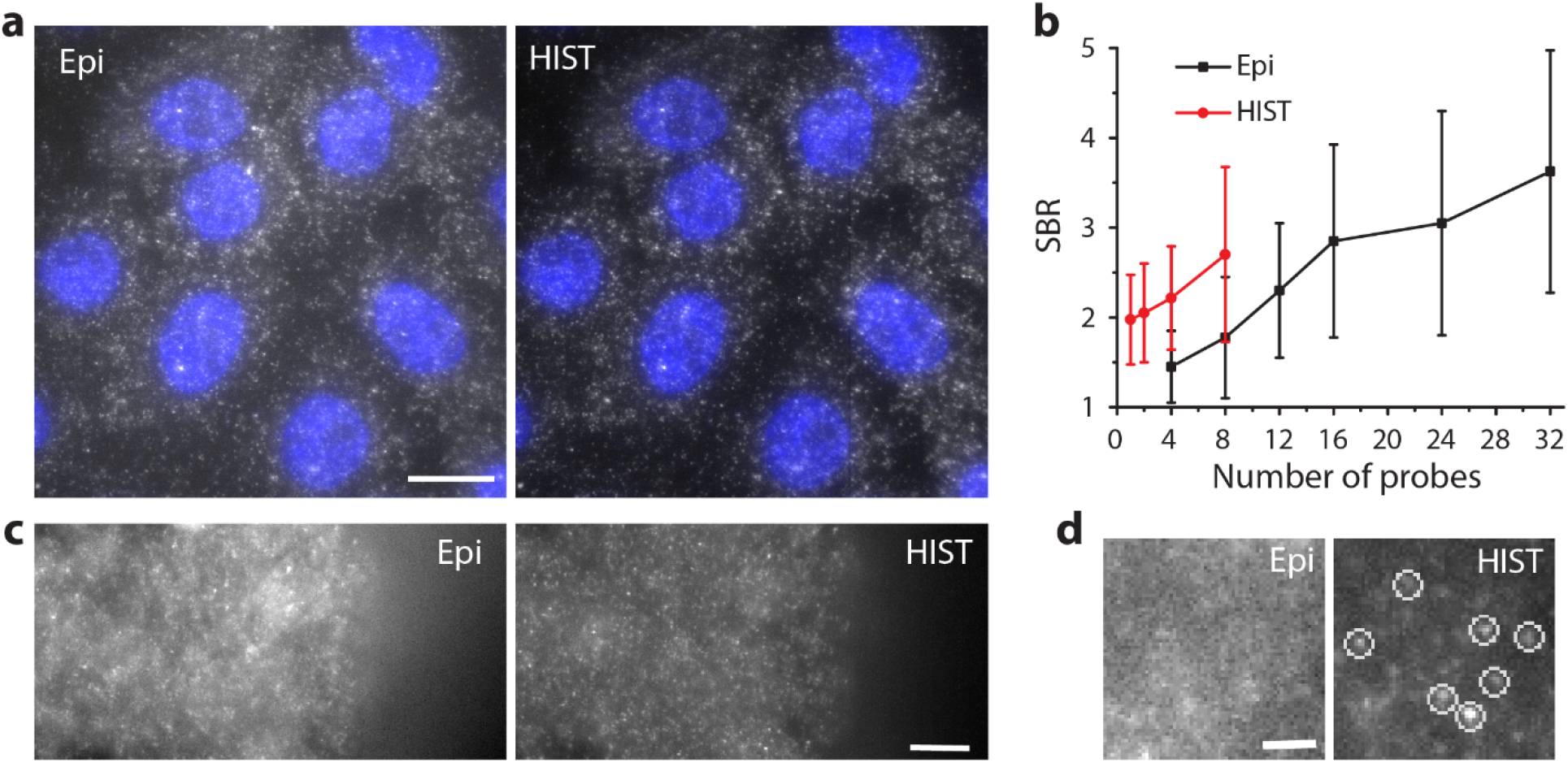
Single-molecule RNA FISH imaging on mammalian cells and tissues with a few probes. (a) smFISH images of *EEF2* on A549 cells with epi and HIST illumination. A maximum intensity projection was performed on 20 z-stacks (5 μm thickness). The illumination power was 40 W/cm^2^ and the integration time was 400 ms. DAPI stain is in blue. (b) Signal to background ratio for different number of FISH probes with epi and HIST illumination. (c) smFISH images of *EEF2* on mouse brain tissue with 5 FISH probes by epi and HIST illumination. The integration time was 800 ms. z-stack images were obtained from *z* = 4 μm to *z* = 7 μm and maximum projected (at coverslip, *z* = 0 μm). (d) Magnified images of the boxed region in (c). White circles indicate identified mRNA spots. All FISH probes were labeled with AF647. Scale bars, 20 μm (a), 10 μm (c), 2 μm (d).

Lastly, we imaged *EEF2* in a 12-μm thick mouse brain tissue with 5 FISH probes. A 3-μm thick z-stack of images was acquired at 5.5 μm in depth. It was difficult to distinguish each individual spot with epi-illumination due to high background while HIST microscopy allowed us to readily detect mRNA (Fig. 2c,d), where SBR was 1.4 ± 0.5. A control experiment with RNase treatment ensured that our FISH probes bound to the target specifically (Supplementary Fig. 10). Note that a standard protocol for smFISH in tissues usually requires at least 48 probes unless the signal is amplified^18^.

Our HIST imaging system provides thinner and much wider illumination compared to HILO imaging, and can be easily implemented onto a standard inverted microscope. It also features much higher photon collection efficiency compared to oblique illumination imaging^19^. It will be feasible to obtain much thinner illumination by using a larger compression ratio. We anticipate that HIST microscopy will benefit several applications including super-resolution imaging, single-molecule tracking and smFISH-based high-throughput gene expression profiling.

## Methods

### HIST microscope

All images were acquired by our custom-made microscope. Three lasers (405 nm, 561 nm and 638 nm; Cobolt) were coupled to a single mode fiber (Thorlabs) and their powers were controlled by a combination of a polarizing beam splitter and a half-wave plate. The fiber output was collimated by a lens (L1, f = 80 mm) and sent to a telescope composed of two cylindrical lenses (CL1, f = 400/250 mm; CL2, f = 50 mm) to generate a tile beam compressed 8x or 5x. The beam was relayed by another telescope system (L2, f = 60 mm; L3, f = 150 mm) with a single-axis galvo mirror (GVS211, Thorlabs), passed through a lens (L4, f = 400 mm), reflected by a dichroic mirror (Di03-R405/488/561/635-t3, Semrock) and focused onto the back focal plane (BFP) of an objective (PlanApo, 60x/1.45, Olympus). The galvo mirror conjugated to the back focal plane was used to sweep the illumination beam on the imaging plane, and controlled by a function generator (DG1032Z, Rigol). A beam incidence angle (*θ*) was adjusted by a mirror (M) conjugated to the imaging plane. A 3-axis piezo stage (MAX311D, Thorlabs) was used for holding samples and acquiring z-stack images, which was controlled by an analog output board (PCI-6733, National Instruments). Fluorescence emission was collected by the same objective and passed through a filter (FF01–446/523/600/677, Semrock). The emission light was then focused on a sCMOS camera (ORCA-Flash4.0 LT, C11440–22C, Hamamatsu) by a tube lens (f = 300 mm). Optionally, a 1:1 4*f* relay system (f = 100 mm) with a slit at the conjugated imaging plane can be inserted before the tube lens to reduce additional scattered light. The function generator sent a master trigger signal to the camera and piezo stage for synchronizing the image acquisition. The sCMOS camera was run in external rolling shutter trigger mode with a line integration time of 60 ms and a delay time of 0.36 ms per line, which corresponds to 800 ms per frame. See Supplementary Fig. 2 for details.

### Single-molecule imaging on 3D hydrogel

A hydrogel solution was prepared using 7.5% acrylamide:bisacrylamide (29:1) (National Diagnostics), 0.2% (v/v) TEMED and 0.02% (w/v) ammonium persulfate in 0.75x TAE buffer. An 18-nt single-stranded DNA with an acrydite moiety at 5’ and Atto647N at 3’ end (called probe 1) was added to the hydrogel solution at a final concentration of 4 nM. 50 μL of this mixture was dropped on a clean coverslip, sandwiched with another coverslip, and incubated for 1 hour at room temperature in the dark. After gently separating them, a thin hydrogel layer about 40 μm thick was washed with and incubated in TAE buffer for 1 hour to remove unbound DNAs. We used an imaging buffer composed of 1x TAE, 0.8% (w/v) dextrose, 1 mg/mL glucose oxidase, 0.04 mg/mL catalase and 2 mM Trolox during all experiments^20^. All chemicals and DNA were purchased from Sigma-Aldrich and IDT unless specified. Please see Supplementary Table 1 for complete sequence information of oligos.

### Imaging analysis

All images were 2×2 binned and the pixel size was 130 nm. When obtaining background subtracted images, a local minimum value in a sub-region (16×16 pixels) was calculated and smoothed to obtain a background image which was subtracted from the raw image with an offset to avoid a negative value. For the calculation of signal-to-background ratio (SBR), the fluorescence images with different illumination methods were acquired at an imaging depth of ~5 μm. The fluorescence intensity of each isolated imaging spot was summed in a 7×7 pixel area around the peak. The corresponding background level (I_B_), which was averaged from the surrounding pixels of each selected spot, was used for fluorescence intensity correction. For all the calibrations, SBRs were defined as: SBR = I_S_/I_B_, where I_S_ was the sum of the fluorescence intensity of central pixels around the peak divided by the average spot size (from point spread function measurement). In each illumination case, more than 100 independent detectable imaging spots were used for S/B comparison.

### Fluorescent nanoparticle imaging on 3D hydrogel

Similar to previous experiments, 20 nm diameter crimson beads (ThermoFisher, F8782) were mixed with a 12% hydrogel solution. 50 μL of the mixture was injected into a flow chamber, and after 10 min the image was measured by an industrial CMOS camera (DMK 33UX290, The Imaging Source) with 200 mm focal length tube lens.

### Characterization of HIST microscopy

For the beam thickness measurement, 200 nm beads (ThermoFisher, F8806) embedded in 7.5% hydrogel were used. Upon a tile beam illumination with 638 nm or 561 nm laser, fluorescence intensities of each bead at different detection planes (z’) were measured by moving a detector with a micrometer. We obtained a corrected depth (z) considering a longitudinal translation of the sample plane and that of the detector plane which are related as z’/z = (1/*n*)×*M*^2^ where *n* is a refractive index of the sample and *M* is a transverse magnification of the system. The intensity at each depth z was averaged from >50 beads and the full width at half maximum (FWHM) of the intensity profile was used for estimating the beam thickness. For the effective width measurement, we illuminated a tile beam with a compression ratio of 5 or 8 on 20 nm bead hydrogel sample, and carried out a standard-deviation y-projection^21^. It results in a line profile along the x-axis, and FWHM of the profile was used as the beam width.

### Photobleaching

To compare photobleaching rates of epi and HIST illumination, the probe 1 in a hydrogel was imaged over 130 μm x 130 μm x 5 μm imaging volume. The exposure time of each frame was 800 ms and an average laser power measured on the back focal plane was 8 mW for both illuminations. The fluorescence intensity of the z-stack was summed and plotted over time. The decay rate of fluorescence intensity was obtained by a single exponential fit to the time trace.

### smFISH on cultured mammalian cells

A549 cells (human lung carcinoma, ATCC CCL-185) were cultured with F-12K medium (ATCC, 30–2004) supplemented with 10% fetal bovine serum (F2442, Sigma) and 1% penicillin/streptomycin (ThermoFisher, 15140122). They were plated on an 8-well Lab-Tek chamber and incubated at 37°C with 5% CO_2_ for 48–72 hours. Cells were fixed with 4% (v/v) paraformaldehyde (PFA; Electron Microscopy Sciences, 15710) at room temperature for 10 min. After washing three times with 1× PBS, cells were permeabilized by 0.5% (v/v) Triton X-100 in 1× PBS for 15 min. After again washing three times with 1× PBS, cells were incubated overnight at 37°C with a various number of FISH probes (*N* = 32, 24, 16, 12, 8, 4, 2, 1) in a hybridization buffer (100 mg/ml dextran sulfate, 1 mg/ml *E.coli* tRNA (Roche, 10109541001), 2 mM Vanadyl ribonucleoside complex (New England Biolabs, S1402S), 0.2 mg/ml RNase free bovine serum albumin (Ambion, AM2616), 2× SSC, 10% deionized formamide (Ambion, AM9342)). The concentration of each probe used was 2.5 nM. The probes against *EEF2* were designed by Stellaris Probe Designer, and 5’ amine modified probes were purchased from IDT. All the *EEF2* probes were labelled with AlexaFluor647 (AF647, Invitrogen, A10277). The nuclei were stained with DAPI. z-stack images were obtained at 0.25 μm steps with the illumination intensity of 40 W/cm^2^. After maximum projection of the z-stack images, the SBR was calculated as described above. Additionally, we measured the photobleaching steps of 4 FISH probes with an illumination power of 150 W/cm^2^ at a fixed imaging plane. At least more than 200 time-traces were used for constructing the distribution of photobleaching step.

### smFISH on mouse brain tissues

Wild-type C57BL6 male mice were used in the present study. The mice were transcardially perfused with 1x PBS containing heparin (10 units/mL) and fixed with 4% PFA in 0.1 M phosphate buffer (pH 7.4). Mouse brains were recovered, post-fixed in 4% PFA solution overnight at 4°C and kept in 30 % sucrose in 1× PBS at 4°C until sinking. The brain was coronally sectioned with 12 μm thickness using a cryostat, collected in 1× PBS. The brain tissue section was gently transferred to an 8-well Lab-Tek chamber supplemented with 500 μL 1× PBS. The tissue was permeabilized with 200 μL of 0.5% Triton X-100 for 25 min at room temperature. After washing out three times with 1× PBS, the brain tissue was stained with 5 FISH probes labeled with AF647 overnight in the hybridization buffer at 37°C. The sample was rinsed with wash buffer (10% deionized formamide in 2× SSC), incubated in 2× SSC for an hour at 37 °C and supplemented with imaging buffer before imaging. For the nonspecific binding test of FISH probes, we prepared the tissue sample like above except that 0.5% RNase A (TheromoFisher, 12091021) was added in the hybridization buffer. All work with mice was performed in accordance with Institutional Animal Care and Use Committee of the University of Central Florida.

### Data availability

The data sets generated in this study are available from the corresponding author upon reasonable request.

## Acknowledgments

We thank Goun Je for preparation of brain tissues, and Hunt Optics for generously loaning the sCMOS camera. This work was supported by start-up funds from CREOL at University of Central Florida.

## Author contributions

J.T. prepared the samples, performed the experiments and analyzed the data. K.Y.H. conceived, guided and supervised the project. J.T. and K.Y.H. wrote the manuscript.

## Competing financial interests

University of Central Florida has filed a patent application covering the work described in this paper.

